# Rapid assay optimization by plug-and-play MAPPIT

**DOI:** 10.1101/2025.09.05.674518

**Authors:** Stefanie Maes, Louis Delhaye, Robin Cooreman, Eline Soetens, Annick Verhee, Delphine De Sutter, Tessa Van de Steene, Long Nguyen, Dominique Audenaert, Xavier Saelens, Sven Eyckerman

**Affiliations:** VIB-UGent Center for Medical Biotechnology, VIB, Ghent, 9052, Belgium; Department of Biomolecular Medicine, Ghent University, Ghent, 9000, Belgium; Department of Biochemistry and Microbiology, Ghent University, Ghent, 9000, Belgium; VIB Screening Core, VIB, Ghent, 9052, Belgium; Centre for Bioassay Development and Screening (C-BIOS), Ghent University, Ghent, 9052, Belgium

**Author notes:** These authors contributed equally (Stefanie Maes, Louis Delhaye).

**Keywords:** MAPPIT, protein-protein interaction, Golden Gate cloning, VHH, MXA

## Abstract

Studying protein-protein interactions, which are crucial for all cellular processes in both health and disease, offers valuable insights into underlying mechanisms and reveals promising therapeutic targets. While efforts have been made to vastly increase the scale of MAPPIT assays, covering the full protein interactome remains challenging, in particular when considering the availability of different assay configurations. In this study, we expanded our Golden Gate modular cloning platform to enable rapid, customizable construct assembly for MAPPIT assays. We demonstrate the strength of this expanded toolbox for multiple distinct protein-protein interactions, including a MAPPIT assay for six different intracellular VHH targeting the anti-viral effector protein MXA.

## Introduction

Protein-protein interactions (PPI) are crucial for all biochemical and regulatory processes that occur in cells. Identifying and characterizing PPIs has been an ongoing effort for understanding disease and immune defense mechanisms^1-3^, as well as for the development of PPI-modulating drugs for various clinical applications^4-7^. Traditional PPI studies focus on a particular protein-of-interest (POI) for which PPI partners are identified through a single PPI discovery method, followed by orthogonal validation using a complementary PPI assay. However, due to method-inherent technical limitations and biological complexity of the research question, a single method is unable to cover the full interactome of any given POI^8-12^.

Methods for PPI discovery are typically classified into binary and mass spectrometry-based approaches^13^. The latter includes techniques such affinity purification coupled to mass spectrometry and proximity labeling, providing proteome-wide coverage and enabling the identification of both direct interaction partners and multi-protein assemblies. However, due to the cost and inherent limitations of current mass spectrometers, high-throughput screening of multiple POIs in parallel remains a challenging endeavor. Despite ongoing advancements, proximity labeling studies have so far examined only a few hundred POIs^14-17^. Similarly, to date, a limited set of seminal affinity purification studies have achieved large-scale coverage, with profiling of roughly 1,000 to 10,000 POIs^18-21^ in HEK293T, HeLa, or HCT116 cells. In contrast, binary methods enable scalable screening of direct PPIs in parallel at nearly proteome-wide scale. For example, Luck *et al*.^22^ performed pairwise screening of 17,408 human proteins with three yeast two-hybrid (Y2H) assay versions.

Binary methods co-express two candidate interacting proteins, a bait and a prey protein, in a cellular system with PPI-dependent activation of a reporter gene serving as a quantifiable readout. Y2H, the first binary high-throughput PPI discovery assay, is based on the PPI-dependent reconstitution of a transcription factor in *Saccharomyces cerevisiae*. Ever since, the core idea has been expanded to incorporate other host organisms and reporter assays^13^. The mammalian protein-protein interaction trap (MAPPIT) is a binary method in mammalian cells which leverages PPI-dependent complementation by decoupling the type I cytokine receptor signal transduction pathway^23^. Type I cytokine receptors constitutively interact with cytosolic JAK kinases. Ligand-dependent receptor clustering triggers the transphosphorylation of JAK kinases, facilitated by the proximity of their kinase domains, which results in the activation of downstream STAT signaling. In MAPPIT, a bait protein is fused to the C-terminus of a chimeric receptor comprising the extracellular domain of the human erythropoietin receptor (EpoR), and the transmembrane and intracellular domains of a JAK2-STAT3 signaling-deficient mutant of the murine leptin receptor (LepR). Prey proteins are fused to the C-terminal fragment (amino acids 760-918) of human gp130 (IL6ST), which can restore downstream JAK2-STAT3 signaling upon bait-prey interaction in a ligand-dependent manner. A STAT3-responsive reporter gene produces a quantifiable signal. Applications of MAPPIT include but are not limited to the validation of large-scale PPI networks^24^, screening of cDNA libraries for novel PPIs^23, 25^, and mapping PPI interfaces by combining with mutagenesis^26^. Ever since, the original MAPPIT platform has been expanded with several variations of the original assay, enabling detection of: modification-dependent interactions (heteromeric MAPPIT^27^), PPI modulators (reverse MAPPIT^28^), protein targets of small molecules (MASPIT^29^), and a βc-MAPPIT^30^ variant developed in hematopoietic cells. This demonstrates that MAPPIT is a versatile binary platform easily adaptable to various research objectives while also allowing scalability.

While efforts to perform MAPPIT screens for specific bait proteins to proteome-scale ORF collections (14,816 prey proteins) have been performed^25^, performing MAPPIT assays in a proteome-by-proteome setup remains challenging. At most, a 2,000-by-2,000 MAPPIT test space has been performed as a validation strategy for Y2H screening^22^. To further increase MAPPIT’s test space, we expanded our Golden Gate modular cloning platform^31^ to include customizable assembly of both MAPPIT bait and prey constructs, improving cloning feasibility. This integration enables alignment of MS-based discovery with orthogonal validation of the MAPPIT assay. In addition, we made an effort towards a standardized MAPPIT setup. Ever since the original MAPPIT publication, alternatives to the original hEpoR-mLepR chimeric receptor configuration have been developed^26, 30, 32, 33^. To date, a comparative experiment to determine whether there is an optimal chimeric receptor configuration with the highest signal-to-noise ratio independent of the fused bait has not been reported. As a consequence, any new MAPPIT experiment requires either an arbitrary choice based on previous literature or a bait-specific comparative experiment that includes time-consuming cloning steps. We used the novel MAPPIT toolbox within our Golden Gate modular cloning platform^31^ to rapidly identify an optimal chimeric receptor configuration. For this, we focused on the BCL2 and BIM^34, 35^, and GFP and VHH-GFP4^36-38^ PPIs. Based on MAPPIT assays for both interaction pairs, we propose a standardized MAPPIT setup with an option for rapid optimization when the bait deviates from this standard using the MAPPIT-compatible Golden Gate assembly platform. As a proof-of-concept, this standardized setup was used in MAPPIT assays for six intracellular single domain antibodies (VHHs) targeting human Myxovirus resistance protein A (MXA), an anti-viral effector protein. These results show that this MAPPIT implementation can be an alternative to co-immunoprecipitations to assess intracellular antigen recognition.

## Experimental Section

### Cell lines and tissue culture

Human HEK293T cells^39^ were cultured in DMEM supplemented with 10% FBS. Parental cell lines were maintained in antibiotic-free conditions, transfection experiments were performed with 30 U/mL Penicillin-Streptomycin. Cells were kept under 60-70% confluency and passaged twice a week. Cell lines were confirmed mycoplasma-free by a mycoplasma PCR detection kit. Cells were maintained in a humidified incubator with 5% CO_2_ at 37°C.

### Plasmids and molecular cloning for MAPPIT

The STAT3-responsive pXP2d2-rPAP1-luci reporter^40^ was used for all MAPPIT assays. All MAPPIT bait and prey constructs were generated using our Golden Gate modular cloning platform^31^. Chimeric receptor (bait) and gp130 (prey) fragments, with internal BsaI and BsmBI removed for Golden Gate assembly, as Level-0 modules were generated by PCR, DNA synthesis, or a combination of both. For the chimeric receptor (bait) fragments: B-EpoR-LepR(F3)-C was generated using DNA synthesis, B-EpoR-LepR(Box1)-(GGS)_60_-C was generated by a combination of DNA synthesis (EpoR) and PCR-based amplification of LepR(Box1)-(GGS)_60_ using pSEL(+2L)60GGS-Y480^33^ as DNA template, B-LepR(F3)-C was generated by a combination of DNA synthesis for the intracellular LepR(F3) domains and PCR-based amplification of the extracellular LepR domains using pcDNA5/FRT-LepR-LepR(F3)-CIS^23^ as a DNA template, B-LepR(Box1)-(GGS)_60_-C was generated by PCR-based amplification of the intracellular LepR(Box1)-(GGS)_60_ domains using pSEL(+2L)60GGS-Y480^33^ as DNA template combined with the PCR-amplified fragment of the extracellular LepR domains using pCLG^41^ as DNA template. Partial fragments were combined and amplified using overlap extension PCR^42^ with B and C overhang compatible overhangs in the forward and reverse primer, respectively. In addition to the B overhangs, the forward primer of the final PCR step included a Kozak sequence and start codon in frame with coding sequence of the downstream ORF. For the gp130 (prey) fragments: B-FLAG-gp130-C was generated using PCR-based amplification of FLAG-gp130 using pMG1^23^ as DNA template. PCR forward and reverse primers contained the B and C overhangs, respectively. In addition, a Kozak sequence and start codon were included in the forward primer. E-gp130-FLAG-F was generated by PCR amplification of gp130-FLAG using pMG1C^43^ as DNA template. PCR forward and reverse primers contained E and F overhangs, respectively. In addition, the reverse primer included a stop codon. Full-length MAPPIT bait and prey fragments were cloned into pGGBC or pGGEF Level-0 backbone vectors (GreenGate Cloning System, Addgene kit #1000000036)^44^ depending on the overhangs using BsaI digestion and T4 ligation at a 3:1 (insert:vector) w:w ratio with a minimum of 50 ng vector. Vectors were chemically transformed in ccdB-sensitive DH10B competent cells. Clones were selected with carbenicillin (50 μg mL^-1^), and screened by a diagnostic restriction digest. Positive clones were verified by Sanger sequencing between the BsaI recognition sites.

Level-1 MAPPIT bait and prey transcriptional units (TU) were generated using BsaI-based Golden Gate assembly. Modular cloning was performed as described previously^31^ with the following adjustments: each reaction mixture consisted of 25 ng per Level-0 module and Level-1 backbone, 100 U T4 DNA ligase, and 5 U BsaI-HFv2 in a total volume of 3.75 μL. DNA mixes containing all Level-0 modules and Level-1 backbone were generated by acoustic mixing on an Echo 650 liquid handler. After BsaI/T4 DNA ligase cycling, assembly reactions were chemically transformed in ccdB-sensitive DH10B competent cells. Clones were selected with kanamycin (50 μg mL^-1^), and screened by a diagnostic restriction digest. Level-1 TUs were directly used for transfection in all MAPPIT assays.

### Immunization, library construction, and panning of MXA VHHs

A llama was injected subcutaneously with 150 μg recombinant wild-type human MXA (a kind gift from Dr. Olivier Daumke, Max Delbrück Center, Berlin, DE) formulated in Gerbu LQ3000 adjuvant for a total of six times at weekly intervals. Immunizations and animal handling were performed according to directive 2010/63/EU of the European parliament for the protection of animals used for scientific purposes, and were approved by the Ethical Committee for Animal Experiments at the Vrije Universiteit Brussel (Brussels, BE) in permit No. 13-601-1. Four days after the last immunization, 100 mL of anti-coagulated blood was collected from which peripheral blood lymphocytes were isolated. Total RNA from peripheral blood lymphocytes was used to prepare cDNA using oligo dT primers. VHH coding sequences were amplified from the cDNA library using a nested PCR. Primers are shown in Table S1. Amplicons were digested with PstI and NotI restriction enzymes and subsequently cloned in the multiple cloning site of the pMECS phagemid vector, downstream of the PselB leader sequence and upstream of the HA and His_6_ affinity tags as well as the pIII phage coat protein for phage display. The VHH library was transformed into electrocompetent *Escherichia coli* TG1 competent cells, which were subsequently infected with VCS M13 helper phages, leading to a VHH displaying phage library. This library was subjected to three rounds of bio-panning using 20 μg of recombinant wild-type human MXA immobilized in a well of a microtiter plate. After blocking, phage particles were added to both the coated and uncoated wells, and the microtiter plate was incubated for 1 h at room temperature. Unbound phage particles were removed, and the wells were washed with 0.1% Tween-20 in PBS (PBST). Bound phage particles were eluted with 14% triethylamine and neutralized with 1 M Tris-HCl (pH 8.0). To initiate a subsequent panning round, the eluted phages were used to re-infect exponentially growing *E. coli* TG1 cells in 2xYT medium containing 1% glucose for 30 min at 37°C, after which VCS M13 helper phages were added and the cells were further incubated for 20 min at 37°C. The culture was then centrifuged and the resulting pellet resuspended in fresh 2xYT medium containing 100 μg mL^-1^ ampicillin and 100 μg mL^-1^ kanamycin. After overnight incubation at 37°C, VHH-displaying phages in the culture supernatant were precipitated with PEG and used for a new round of panning. Between panning rounds, phage enrichment was evaluated by serial dilution from coated and uncoated wells and assessed by the number of overnight transformants.

### Seroconversion of llama serum against MXA and mmMx1

Wells of a microtiter plates were coated with 100 μL recombinant wild-type human MXA or murine Mx1 (mmMx1) at a concentration of 2 ng μL^-1^ in PBS. After overnight incubation at 4°C, plates were washed once with PBST, followed by three washes with distilled water. Plates were blocked for 1 h with 1% BSA in PBS, washed again as before, and incubated with serial dilutions of the llama sera in sample buffer (0.5% BSA and 0.05% Tween-20 in PBS) for 2 h at room temperature. MXA- and mmMx1-specific llama IgG were detected with primary goat ɑ-llama IgG (A160-100A, Bethyl Laboratories, Inc.) and secondary donkey ɑ-goat HRP-coupled (ab97110, Abcam; RRID: AB_10679463) antibodies diluted in sample buffer. Absorbance at 450 nm was measured after supplementation of 50 μL 3,3’,5,5’-tetramethylbenzidine substrate and reactions were stopped by addition of 50 μL 1 M H_2_SO_4_ per well.

### Detection of MXA-specific binders in periplasmic extracts

After panning, individual pMECS transformants were cultured for 5 h at 37°C in 2 mL of TB medium containing 100 ug mL^-^ ampicillin. VHH expression was induced overnight with 1 mM IPTG at 37°C. Bacterial cells were then pelleted, resuspended in 200 μL TES buffer (0.2 M Tris-HCl (pH 8.0), 0.5 M EDTA, and 0.5 M sucrose), and incubated for 30 min at 4°C with shaking. The periplasmic content was released by adding 300 μL water to induce an osmotic shock. After 1 h incubation at 4°C, periplasmic extracts were collected by centrifugation. VHH-containing periplasmic extracts were evaluated for MXA binding-specificity by ELISA. A microtiter plate was coated overnight with either 100 ng recombinant human wild-type MXA or 100 ng BSA at 4°C. On the next day, the plates were blocked with 4% milk-powder in PBS, and 100 μL periplasmic extract was added to both the MXA- and BSA-coated wells. Bound VHHs were detected with 1:2000 mouse ɑ-HA (901533, BioLegend; RRID: AB_2314672) and 1:2000 ɑ-mouse HRP-coupled (NXA931, Cytiva; RRID: AB_772209) primary and secondary antibodies, respectively. Absorbance at 450 nm was measured after adding 50 μL 3,3’,5,5’-tetramethylbenzidine substrate and reactions were stopped by addition of 50 μL 1 M H_2_SO_4_ per well. Periplasmic extracts of which the absorbance of the MXA-coated well was two-fold higher than the BSA-coated well were considered to contain MXA-specific VHHs. Phagemid DNA of the selected colonies was extracted and sequenced with the MP057 primer (Table S1).

### Cloning and production in Pichia pastoris, and Nickel-Nitrilotriacetic acid (Ni-NTA) purification of VHHs

VHH coding sequences were amplified using the pMECS phagemid as a template. Primers are shown in Table S1. PCR fragments were digested with XhoI and SpeI restriction enzymes and ligated into pKai61^45^ in frame with a *Saccharamyces cerevisiae* α-mating factor signal sequence and a His_6_-tag, and under the control of the MeOH-inducible AOX1 promoter. pKai61-VHH plasmids were linearized by PmeI digestion and transformed into *Pichia pastoris* GS115 cells using the condensed transformation protocol^46^. Two to five *Pichia pastoris* transformants were inoculated in 2 mL YPNG medium containing 100 μg mL^-1^ zeocin for 24 h at 28°C. Cells were pelleted, resuspended in 2 mL YPNM medium, incubated at 28°C and spiked with 50 μL of 50% MeOH at 16 h, 24 h, and 40 h of incubation. After 48 h of incubation, cells were pelleted by centrifugation and the supernatant was collected. VHH presence in the supernatant was confirmed by SDS-PAGE and Coomassie Brilliant Blue staining. Clones that yielded high VHH expression in the medium were selected and used for 300 mL productions. Growth and MeOH induction conditions and harvesting of medium were as described above. VHH in the cleared culture medium were precipitated by the addition of (NH_4_)_2_SO_4_ up to 80% saturation, followed by incubation for 4 h at 4°C. Precipitated VHHs were pelleted by centrifugation at 20.000 x g, resuspended in 10 mL HisTrap binding buffer (20 mM (NH_4_)_2_SO_4_, (pH 7.5), 0.5 M NaCl, 20 mM imidazole (pH 7.4)), and purified on a 1 mL HisTrap HP column following manufacturer’s instructions. VHH-containing fractions were pooled and concentrated with an Amicon ultra column (3 kDa cutoff) followed by buffer exchange to PBS. The purity of the VHHs was verified by SDS-PAGE and Coomassie Brilliant Blue staining.

### In vitro binding of MXA VHHs to wild-type and M527D MXA

Microtiter plates were coated with 100 ng wild-type or M527D MXA after which recombinant VHHs were added in a four-fold serial dilution starting at 80 μg mL^-1^. VHH binding to MXA was detected using 1:2000 mouse ɑ-His_6_ (MCA1396GA, Bio-Rad; RRID: AB_566361) and 1:2000 ɑ-mouse HRP-coupled (NXA931, Cytiva; RRID: AB_772209) primary and secondary antibodies, respectively. Absorbance was measured as described above. Curve fitting was performed by dose-response analysis in R using a four-parameter log-logistic function (LL.4) implemented in the drc package.

### Prey expression profiling

To assess prey expression, 34,000 HEK293T (passage < 10) cells were seeded in a 48-well plate. The next day, cells were transfected using Ca_3_(PO_4_)_2_. DNA mixes containing 275 ng prey Level-1 plasmid and 250 mM CaCl_2_ in 17 μL total volume, were mixed dropwise with 17 μL 2X BioUltra HEPES-buffered saline (HEBS) solution (Sigma-Aldrich) while vortexing. Transfection mixtures were incubated 10 min at room temperature, and transferred onto the cells without disturbing them. The next day, cells were washed once with PBS, and lysed directly on the plate with 30 μL RIPA lysis buffer (25 mM Tris-HCl (pH 8.0), 75 mM NaCl, 1 mM EDTA, 0.5% NP40, 0.05% SDS, supplemented fresh with 100X cOmplete protease inhibitor cocktail (Roche) and 0.025% (v:v) f.c. sodium deoxycholate). Lysates were scraped on ice, transferred to a microcentrifuge tube, and incubated 15 min on ice to promote lysis. Lysates were cleared by centrifugation at 16,100 x g at 4°C for 15 min. Supernatant was transferred to a new microcentrifuge tube, and stored at -20°C until further processing.

### Co-immunoprecipitation

VHH coding sequences were amplified with the pMECS phagemids as a template using the primers shown in Table S1. Amplicons were digested with XhoI and NotI restriction enzymes and ligated in the multiple cloning site of pCAGGS vectors for mammalian expression. 1.28 X 10^6^ HEK293T cells were seeded in 6-well plates, and, the next day, co-transfected with 500 ng pCAGGS-VHH-His_6_ and 500 ng pCAGGS-FLAG-MXA expression constructs using FuGENE HD according to the manufacturer’s instructions. Twenty-four hours after transfection, cells were washed once with ice-cold PBS and scraped in 250 μL mild lysis buffer (50 mM HEPES (pH 8.0), 100 mM KCl, 2 mM EDTA, 0.1% NP-40, 10% glycerol, supplemented fresh with 100X cOmplete EDTA-free Protease Inhibitor Cocktail (Roche). Cells were incubated on ice for 15 min to promote lysis. Lysates were cleared by centrifugation at 16.000 x g for 15 min at 4°C. Twenty microliters of input control was kept, and the remaining lysate was incubated with 20 μL equilibrated mouse ɑ-FLAG M1 (F3040, Sigma-Aldrich; RRID: AB_439712) for 2 h at 4°C with end-over-end rotation. Beads were washed four times with mild lysis buffer, and bound proteins were eluted by incubation with 20 μL 3X Laemmli buffer at 95°C for 5 min. Eluted protein samples were analyzed by immunoblotting. A non-neutralizing VHH raised against the RSV fusion protein (RSV-F) from the VHH library described in Rossey *et al*.^47^ was used as a negative control VHH.

### Immunoblotting and antibodies

Protein concentration of the lysates was measured using the Bradford method in technical triplicates per lysate, and 30 μg protein was supplemented with 7.5 μL XT Sample Buffer and 1.5 μL XT Reducing Agent in a final volume of 30 μL in water. Samples were heated to 95°C for 10 min, cooled down on ice to room temperature, and loaded on a 4-12% ExpressPlus PAGE pre-cast gel (Genscript) according to the manufacturer’s instructions. The gel was ran for 10 min at 100 V, followed by 1 h at 140 V. Proteins were transferred to PVDF membrane for 30 min at 100 V in methanol blotting buffer (48 mM Tris-HCl, 39 mM glycine, 0.0375% SDS (w:v) and 20% methanol (v:v)). Membranes were blocked at room temperature for 30 min in Odyssey Blocking buffer (LI-COR) with end-to-end rotation. Primary antibodies were incubated overnight at 4°C with gentle end-to-end rotation. Mouse α-FLAG (R960-25, Invitrogen; RRID_2556564), mouse α-His_6_ (MCA1396GA, Bio-Rad; RRID: AB_566361), rabbit α-ACTB (A2066, Sigma-Aldrich; RRID: AB_476693), and rabbit ɑ-GAPDH (ab9485, Abcam; RRID_307275) primary antibodies were diluted in TBS at 1:5000, 1:2000, 1:2000, and 1:2000, respectively. Polyclonal rabbit anti-MXA was commercially generated by immunization of a rabbit at days 0, 14, 28, and 56 with 100 μg wild-type human MXA (CER Groupe, BE). Serum was collected at day 66 and the IgG fraction was enriched using a MabSelect SuRe column (Cytiva) following the manufacturer’s instructions. Membranes were washed three times with TBST for 10 min at room temperature with gentle end-to-end rotation, after which membranes were incubated with secondary antibodies for 1 h at room temperature with end-to-end rotation. Goat polyclonal α-mouse IgG IRDye 800CW (LI-COR) and goat polyclonal α-rabbit IgG IRDye 680RD (LI-COR) were used at a dilution of 1:5000 in TBS. After secondary antibody staining, membranes were washed again three times with TBST, once with TBS, and visualized on a LI-COR Odyssey IR scanner. Uncropped immunoblots are shown in Fig. S3.

### MAPPIT

MAPPIT experiments were performed as described previously^48^. Leptin (LEP) and erythropoietin (EPO) were used at final concentrations of 100 ng mL^-1^ and 4 ng mL^-1^, respectively. Each condition was performed in triplicate using 10,000 HEK293T cells per replicate followed by ligand or solvent supplementation to the growth medium. For ‘mock’ conditions we transfected the pSV-SPORT vector. Fold inductions were calculated as the ratio of stimulated relative light units (RLU) per replicate over the average of all unstimulated RLUs per condition. Statistics were performed in R with the limma package. A linear model was fitted onto the data and differential analysis was performed by empirical Bayes moderated one-sided t-tests with a Benjamini-Hochberg multiple testing correction. Fold changes (FC) were calculated over mock conditions and log2 transformed. Adjusted *P*-values below or equal to a 5% threshold were considered statistically significant. Fold induction of all MAPPIT experiments can be found in Table S2.

### VHH Kabat numbering, multiple sequence alignment, and phylogenetic analysis

The amino acids of the MXA-binding VHHs were numbered according to the Kabat numbering scheme using the language model implemented in ANARCII^49^ with CDRs defined accordingly. VHH sequences were aligned using MAFFT with default BLOSUM62 substitution matrix and gap penalties as implemented in NGPhylogeny.fr^50^. Dendrograms were inferred using the PhyML with automatic Smart Model Selection (SMS) on NGPhylogeny.fr^50^ to choose the best-fitting substitution model. Branch support was estimated by approximate likelihood ratio test (aLRT). VHH-GFP4, VHH-IAV-M2e, and VHH-RSV-F were used as outgroups to re-root the phylogenetic tree.

## Results and Discussion

### Expanding the Golden Gate modular cloning platform for PPI analysis with MAPPIT

We previously developed a Golden Gate modular assembly platform tailored for proximity labeling proteomics using TurboID^31^. Here, we expand this platform with a MAPPIT toolbox. We constructed a set of Level-0 modules encoding four distinct chimeric receptor configurations, based on earlier studies^23, 26, 33^ (Fig. 1A). These chimeric receptor modules consist of combinations of the human erythropoietin receptor (EpoR) or the murine leptin receptor (LepR) extracellular domains, fused to one of two signaling-deficient intracellular LepR variants. The intracellular domains, including the transmembrane region, are based on the long LepR isoform. One variant carries triple tyrosine-to-phenylalanine substitutions (F3; Y987F, Y1077F, and Y1138F) to abrogate downstream JAK-STAT signaling. The second variant consists of the LepR Box1 motif, which is required for JAK2 recruitment, followed by a synthetic tail comprising a 20x Gly-Gly-Ser (GGS) tandem repeat, serving as a flexible spacer. To preserve the endogenous N-terminus of the chimeric receptor constructs, we cloned them into BC Level-0 modules to avoid the Gly-Ser scar sequence inherent to the other positions in the Golden Gate assembly platform (Fig. 1B). Bait proteins were incorporated at the CD position. Similarly, gp130 Level-0 modules for the MAPPIT prey fusion were generated for N- and C-terminal tagging of gp130 (amino acids 760-918 of human IL6ST). To this end, FLAG-gp130 with a C-terminal Gly-Gly-Ser hinge and gp130-FLAG fragments were cloned in the BC and EF Level-0 modules, respectively. Prey proteins were incorporated at the CD position (Fig. 1B).

**Fig. 1:**
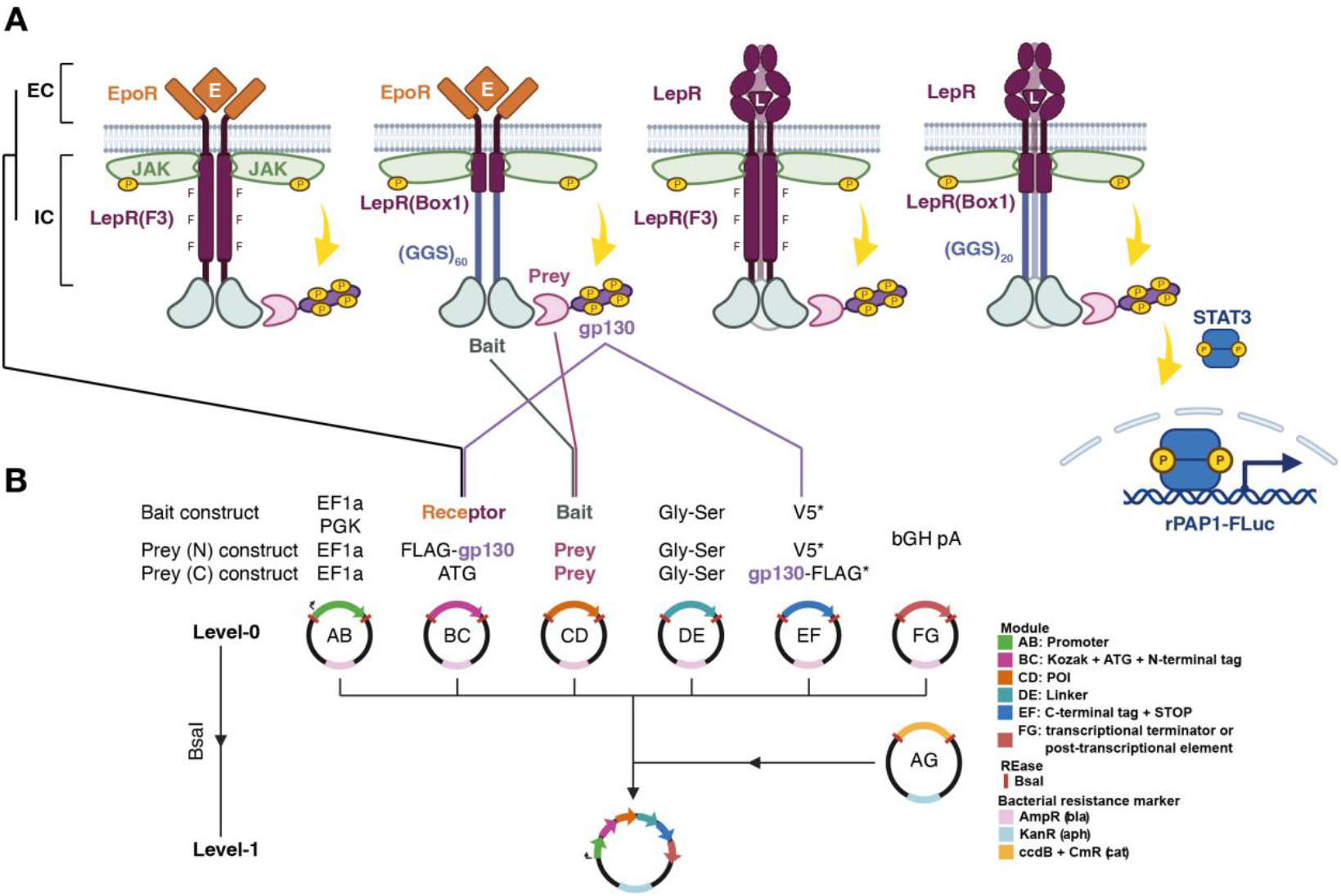
A Golden Gate cloning platform for MAPPIT. (A) Overview of the MAPPIT principle for all four different chimeric receptor bait configurations. (B) Integration of MAPPIT components into a Golden Gate modular cloning platform. BsaI overhangs are shown for each position. EC, extracellular; IC, intracellular; EpoR, erythropoietin receptor; LepR, leptin receptor; POI, protein-of-interest; N, N-terminal; C, C-terminal. P in yellow circles represents phosphorylation events, F represents Y987F, 71077F, and Y1138F mutations, E represents the erythropoietin ligand; L represents the leptin ligand.

### Towards a standardized MAPPIT setup

We used the MAPPIT toolbox for rapid validation of the existence of a general chimeric receptor configuration for the MAPPIT bait. To this end, we performed comparative experiments to identify the chimeric receptor configuration with the highest signal-to-noise ratio independent of the fused bait protein. We focused on two distinct PPIs: the interaction between the apoptosis regulatory proteins BIM (BCL2L11) and BCL2^34, 35^, and the interaction between GFP and VHH-GFP4, a well-characterized GFP-binding VHH^36-38^. Using the Golden Gate modular cloning platform, we assembled bait constructs in all four chimeric receptor configurations under the control of two different promoters as well as N- and C-terminal prey construct for each of all four proteins. Prey expression in HEK293T cells was validated by immunoblotting (Fig. S1A). To confirm that the prey constructs do not auto-activate the STAT3-responsive reporter, we measured reporter activity in the absence of a bait. EpoR-LepR(FFY), a chimeric receptor that retains STAT3 recruitment through phosphorylated Y1138, and the wild-type LepR in the absence of a prey were included to verify ligand-dependent signaling activity. No prey construct showed any significant receptor-independent signaling induction of the reporter compared to mock conditions after EPO or LEP supplementation (Fig. S1B). We continued by performing MAPPIT assays for all bait constructs co-transfected with N-terminal prey constructs (Fig. 2A). In addition to a mock condition, we included a gp130-VHH-GFP4 and gp130-BIM construct as irrelevant prey for the BIM-BCL2 and GFP-VHH-GFP4 interactions, respectively. Except for EF1a-EpoR(Box1)-BIM, assays with irrelevant preys showed no significant induction compared to mock for BCL2 and BIM bait proteins. In contrast, for GFP and VHH-GFP4 various chimeric receptor configurations demonstrated induction when transfected with gp130-BIM. Nonetheless, the fold induction was consistently lower compared to GFP and VHH-GFP4 prey. For chimeric receptor configurations of the BCL2, VHH-GFP4, and GFP baits, a significant reporter induction was observed compared to mock conditions, with EF1a-LepR(Box1) demonstrating the highest induction across all baits (Fig. 2B). Moreover, for this particular configuration, no significant induction was observed when transfected with an irrelevant prey (Fig. 2A, B). BIM bait constructs showed overall lower reporter induction, with only significant inductions for the EF1a-EpoR(Box1) and both EF1a-LepR configurations, compared to mock. Nonetheless, the EF1a-LepR(Box1) configuration demonstrated the highest reporter induction, consistent with the other baits protein. We performed the same experiment with C-terminal prey constructs compared to mock conditions and observed similar results (Fig. S1C). Consistent with the results from the N-terminal preys, BIM bait constructs tended to induce lower reporter induction compared to the other bait proteins, and the EF1a-LepR(Box1) configuration elicited the highest reporter induction for all baits. When co-transfected with irrelevant C-terminal preys, the EF1a-LepR(Box1) configuration induced substantially lower reporter activity compared to its known binding partner for all baits (Fig. S1D), indicating a low rate of false-positive detection in the assay. These results support a standardized MAPPIT setup comprising a bait in the EF1a-LepR(Box1) configuration and an N-terminal prey. In addition, integration of the MAPPIT toolbox in our Golden Gate cloning platform allows the option for rapid optimization for specific bait and/or prey proteins.

**Fig. 2:**
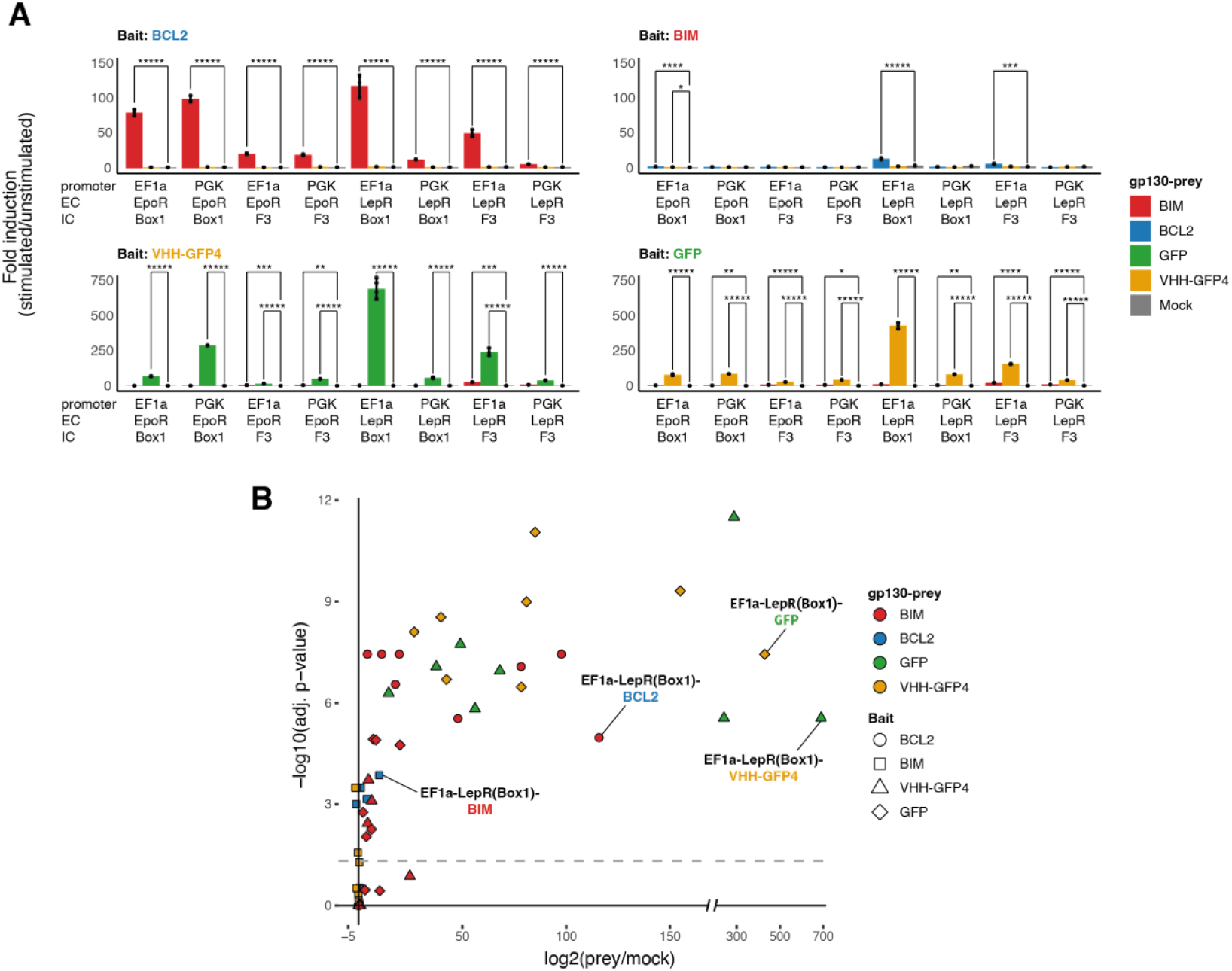
MAPPIT assay optimization using BCL2-BIM and GFP-VHH-GFP4 PPIs. (A) MAPPIT assays for BCL2-BIM and GFP-VHH-GFP4b for all four receptor configurations and both promoters for N-terminal tagged preys (N = 3). (B) Differential analysis (log2(prey/mock)) of all MAPPIT assays (N = 3). The most differential receptor configuration for each MAPPIT assay is highlighted. Dashed line represents adj. P-value of 0.01. EC, extracellular; IC, intracellular; EpoR, erythropoietin receptor; LepR, leptin receptor; F3 represents the Y987F, 71077F, and Y1138F LepR mutant; Box1 represents the Box1 motif LepR mutant with a (GGS)60 cytoplasmic tail. Adj. P-value: * < 5 x 10^-2^; ** < 5 x 10^-3^; *** < 5 x 10^-4^; **** < 5 x 10^-5^; ***** < 5 x 10^-6^.

### MAPPIT as an alternative assay for VHH intracellular antigen recognition

To further explore the standardized assay and show MAPPIT’s utility for studying the binding of intracellular VHH molecules, we performed MAPPIT assays for six MXA VHHs. MXA VHHs were obtained from a VHH phage display library generated from a llama that had been immunized with recombinant human MXA. Antibody specificity was evaluated by testing immune pre-immune serum and immune plasma for reactivity against coated wild-type human MXA or its orthologue wild-type *Mus musculus* Mx1 (mmMx1) (Fig. S2A). MXA immune plasma demonstrated strong immunoglobulin binding to immobilized MXA antigen, with minimal reactivity to immobilized mmMx1. Similarly, immune serum from a llama immunized with mmMx1 did not show reactivity with human MXA, but did with mmMx1 (Fig. S2A). Immune serum of a llama immunized with human metapneumovirus fusion protein (HMPV-F) did not react with either MXA or mmMx1 (Fig. S2A).

The phage display library was subjected to bio-panning on immobilized MXA, leading to a 14,000-fold enrichment of MXA-binding phages after three rounds of panning. A total of 188 bacterial clones obtained after the third panning round were screened for MXA specificity. VHHs were harvested from the periplasm by osmotic shock, and binding of VHHs from periplasmic extracts to MXA and mmMx1 was assessed by ELISA (Fig. S2B). We tested 188 clones over two different microtiter plates; 178 (94.7%) were positive for MXA binding and none demonstrated binding activity to mmMx1. Thirty MXA-positive clones were randomly selected for sequencing, of which 21 were successfully cloned in the *Pichia pastoris* pKai61 expression vectors. Of these, 11 VHHs were selected for further characterization based on their sequence diversity (Fig. S2C-D). These MXA-specific VHHs all contained the conserved cysteine bridge linking FR2 and CDR3 (Fig. S2C), and could be grouped in five different clusters based on sequence similarity (Fig. S2D).

VHHs were expressed in *Pichia pastoris*, and secreted His_6_-tagged VHHs were purified from the culture supernatant by Ni-NTA affinity chromatography. *In vitro* binding of the purified VHHs to wild-type MXA and the monomeric M527D MXA mutant was evaluated by ELISA (Fig. 3A). A negative control VHH targeting the extracellular domain of influenza A virus matrix protein 2 (IAV-M2e)^51^ did not show reactivity with either MXA variant. With the exception of VHH5 and VHH7, most VHHs exhibited a low affinity for wild-type MXA. In contrast, nearly all VHHs, except VHH37 and VHH108, demonstrated a high affinity for the monomeric M527D MXA mutant. Next, we assessed the capacity of the selected VHHs to target MXA intracellularly. All VHHs demonstrated intracellular expression 24 h after transfection of VHH-encoding plasmids in HEK293T cells (Fig. 3B). To determine intracellular binding of these VHHs to MXA, we co-transfected a FLAG-MXA construct with the His_6_-tagged VHH constructs to perform co-immunoprecipitation using an ɑ-FLAG antibody. A negative control VHH targeting the human respiratory syncytial virus fusion protein (RSV-F) was not enriched after MXA pulldown. In contrast, all MXA-targeting VHHs were enriched after FLAG pulldown, demonstrating specific MXA-binding at least in cellular lysates. To verify the VHH-MXA interaction in intact cells, we performed MAPPIT screen with the VHHs or MXA either as prey or bait with the optimized LepR(Box1) receptor configuration under control of the EF1a promoter (Fig. 3C). Six VHHs (VHH4, VHH5, VHH7, VHH29, VHH37, and VHH60) and wild-type human MXA were cloned in the modular cloning platform as outlined before. As control VHH and target, we incorporated VHH-GFP4 and GFP from our proof-of-concept experiments which were previously shown to interact intracellularly^36^. Consistent with our previous experiments, reporter activation was observed only when GFP or VHH-GFP4 bait constructs were co-transfected with their corresponding prey constructs (VHH-GFP4 or GFP, respectively), and not with any of the VHHs targeting MXA. Similarly, no reporter activity was observed for MXA bait or prey constructs co-transfected with VHH-GFP4. In contrast, VHH-MXA bait and prey constructs did demonstrate reporter activation after co-transfection with MXA prey or bait constructs, respectively. These data demonstrate that MAPPIT is capable of screening intracellular VHHs for antigen recognition in an *in cellulo* context with high specificity and no off-target reporter activation.

**Fig. 3:**
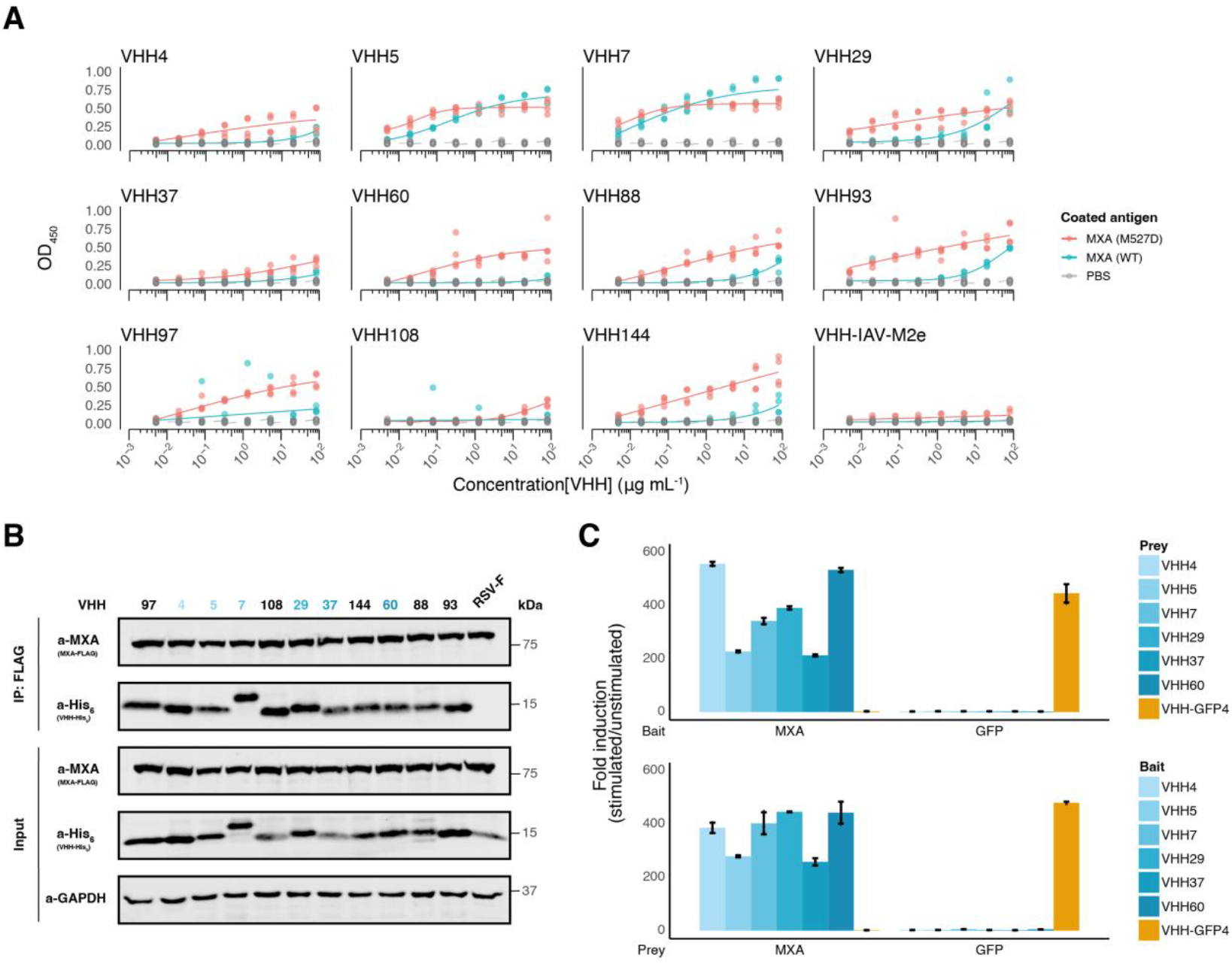
MAPPIT as an alternative to co-immunoprecipitations for intracellular antigen recognition. (A) In vitro binding of eleven selected VHHs and VHH-IAV-M2e to coated wild-type human MXA and M527D MXA (N = 4). (B) Co-immunoprecipitation of intracellularly expressed VHHs after MXA pulldown. Colors of the annotation are in line with panel C. (C) MAPPIT assays for MXA and GFP, and the intracellularly expressed VHH, as either bait or prey constructs (N = 3). VHH-GFP4 is shown as control VHH.

## Conclusions

In summary, we expanded our Golden Gate modular cloning platform with a MAPPIT toolbox that enables rapid assembly of constructs with the option to choose from different setups. This integration in a fully modular cloning system enables MS-based PPI discovery and orthogonal validation in a singular kit. Moreover, it also opens the way to pooled assembly approaches and pooled MAPPIT screening. EF1a-LepR(Box1) was identified as a general optimized setup in a comparative experiment involving two distinct interactions (BCL2-BIM and GFP-VHH-GFP4). This standardized setup can easily be employed as a default for any new MAPPIT assays, as demonstrated for the VHH-MXA intrabodies. In addition, the platform allows rapid optimization of any bait-prey interaction using the alternative available setups. Fully automated modular cloning and tissue culture workflows can even further increase the complexity of MAPPIT assays and further reduce the hands-on time required to perform them, indicating the potential for scaling up to proteome-by-proteome-wide screening.

## AUTHOR INFORMATION

## Corresponding Author

Correspondence should be addressed to

## Present Addresses

† Eline Soetens; Department of Pharmaceutics, Ghent University, Ghent, 9000, Belgium

## Author Contributions

The manuscript was written through contributions of all authors. All authors have given approval to the final version of the manuscript. ^#^ These authors contributed equally.

Contribution Roles Taxanomy (CRediT):

Conceptualization: S.M., L.D., R.C. E.S., X.S., S.E.; Formal analysis: S.M., L.D., R.C., E.S.; Funding acquisition: X.S., S.E.; Investigation: T.v.d.S, D.D.S, A.V., L.N., E.S.; Methodology: S.M., L.D., R.C., E.S., T.v.d.S., D.D.S., A.V., S.E; Project administration: S.M., L.D., D.A., X.S., S.E.; Software: L.D.; Resources: L.N., A.D., X.S.; Supervision: D.A., X.S., S.E.; Visualization: S.M., L.D.; Writing – original draft: S.M., L.D., R.C., S.E.; Writing – review & editing: all authors.

## Notes

The authors declare no competing interests.

## ACKNOWLEDGMENT

The authors would like to acknowledge the VIB Screening Core and VIB Nanobody® VHH Core for their assistance on the work presented in the manuscript. The authors thank Dr. Oliver Daumke (Max Delbrück Center, Berlin, Germany) for providing purified recombinant wild-type and M527D mutant MXA.

## ABBREVIATIONS

PPI: protein-protein interactions
POI: protein-of-interest
Y2H: yeast two-hybrid
MAPPIT: mammalian protein-protein interaction trap
EpoR: human erythropoietin receptor
LepR: murine leptin receptor
MXA: human Myxovirus resistance protein A
TU: transcriptional unit
PBST: 0.1%Tween-20 in PBS
Ni-NTA: Nickel-Nitrilotriacetic acid
HEBS: HEPES-buffered saline
LEP: leptin
EPO: erythropoietin
RLU: relative light units
FC: fold change
SMS: Smart Model Selection
Alrt: approximate likelihood ratio test
mmMx1: *Mus musculus* Mx1
HMPV-F: human metapneumovirus fusion protein
IAV-M2e: influenza A virus matrix protein 2
RSV-F: human respiratory syncytial virus fusion protein

---

**Fig. S1:**
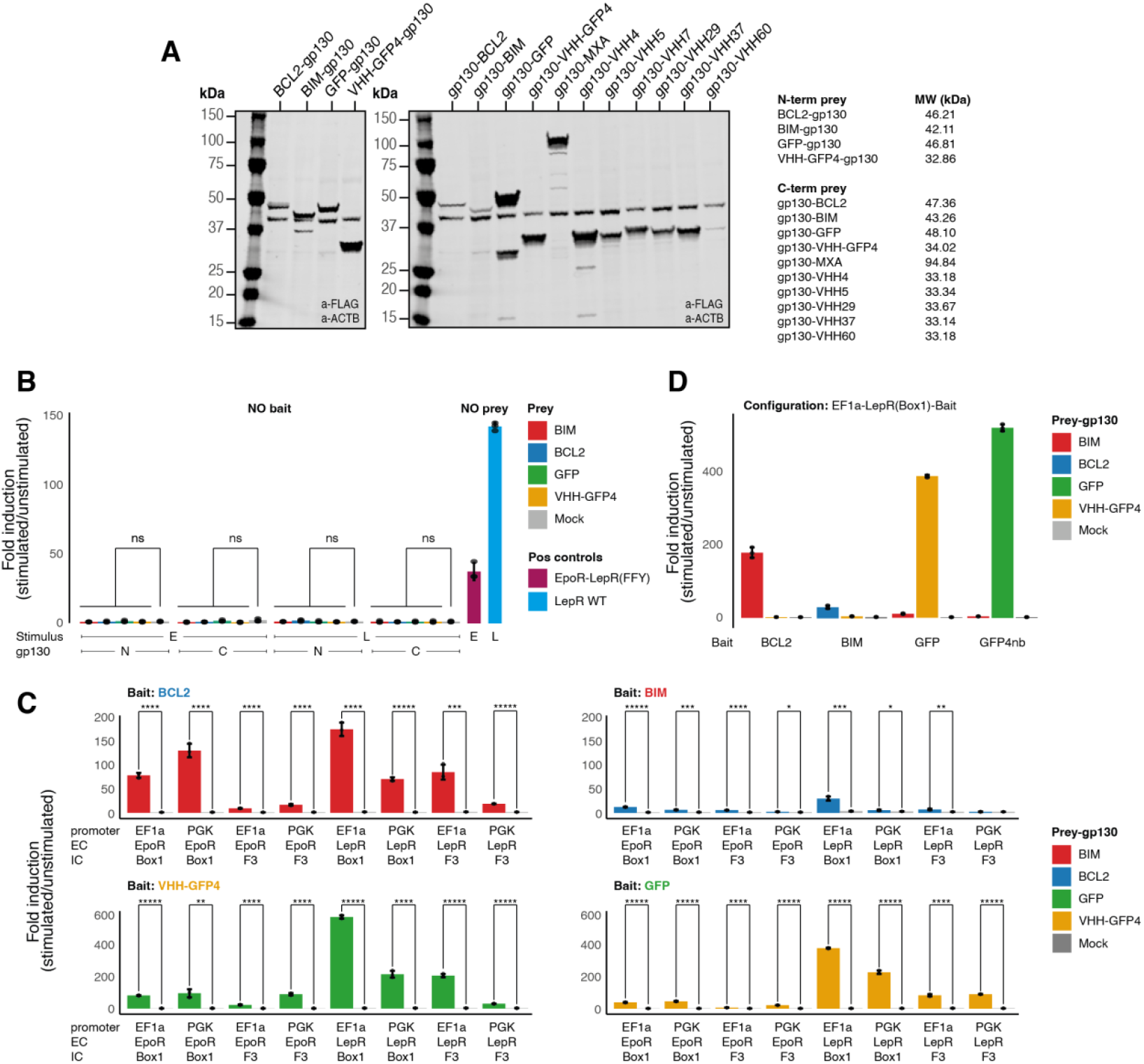
MAPPIT assay optimization using BCL2-BIM and GFP-VHH-GFP4 PPIs. (A) Prey expression assessed by immunoblotting with ɑ-FLAG with ɑ-ACTB as a loading control. Expected molecular weight is shown. (B) Profiling of bait-independent reporter activation after ligand supplementation (N = 3). (C) MAPPIT assays for BCL2-BIM and GFP-VHH-GFP4 for all four receptor configurations and both promoters for C-terminal tagged preys (N = 3). (D) MAPPIT assays of the EF1a-LepR(Box1)-Bait experiments shown in (C) including irrelevant preys (N = 3). MW, molecular weight; EC, extracellular; IC, intracellular; EpoR, erythropoietin receptor; LepR, leptin receptor; F3 represents the Y987F, 71077F, and Y1138F LepR mutant; Box1 represents the Box1 motif LepR mutant with a (GGS)_60_ cytoplasmic tail. Adj. P-value: ns > 5 x 10^-2^; * < 5 x 10^-2^; ** < 5 x 10^-3^; *** < 5 x 10^-4^; **** < 5 x 10^-5^; ***** < 5 x 10^-6^.

**Fig. S2:**
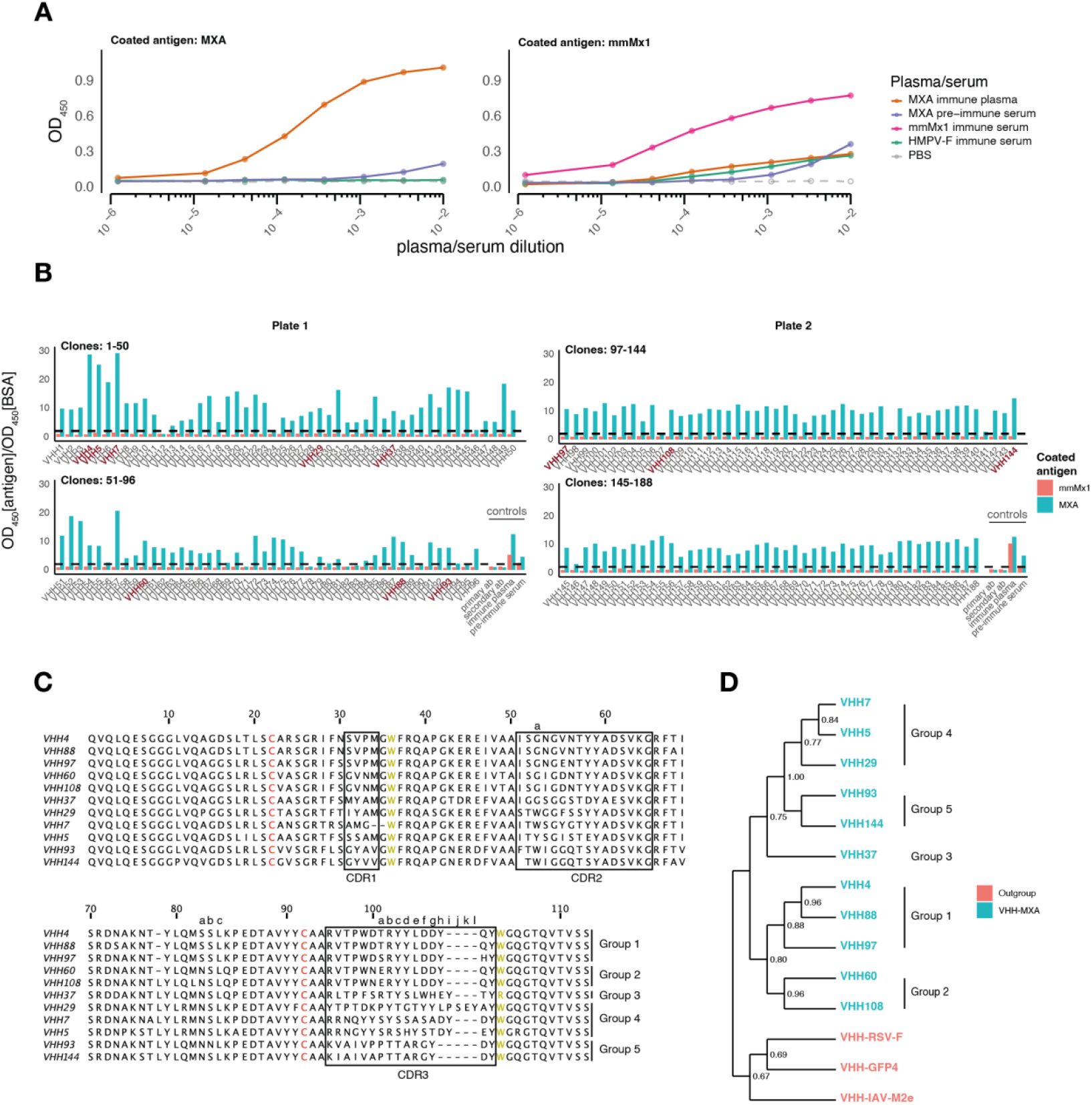
Characterization of a VHH library targeting human MXA. (A) Seroconversion after immunization of a llama with recombinant MXA (N = 1). Murine Mx1 was included to demonstrate specificity of MXA-challenged immune plasma. Sera of llamas immunized with recombinant human metapneumovirus F protein (HMPV-F) or mmMx1 were included as control sera. (B) Screen of E. coli periplasmic extracts of VHH clones isolated after three rounds of bio-panning on human wild-type MXA (N = 1). Primary and secondary antibodies alone were included as technical controls per plate. Immune plasma and pre-immune serum of a MXA-challenged llama was included as biological controls. Dashed line represents the threshold of a two-fold increase over BSA. Selected VHHs are shown in burgundy. (C) Kabat numbering and sequence annotation of selected VHHs. CDR regions are highlighted in boxes, conserved cysteines are shown in red, conserved aromatics are shown in gold. Kabat numbering is outlined above the sequences. Grouping of the sequences was based on phylogenetic analysis after multiple sequence alignment. (D) Dendrogram of the VHHs after multiple sequence alignment. Branch support (aLRT) is shown next to each branching event. The tree was re-rooted to incorporate VHH-RSV-F, VHH-GFP4, and VHH-IAV-M2e as outgroups, as represented by the color code. Grouping was based on supported grouping in the tree.

**Fig. S3:**
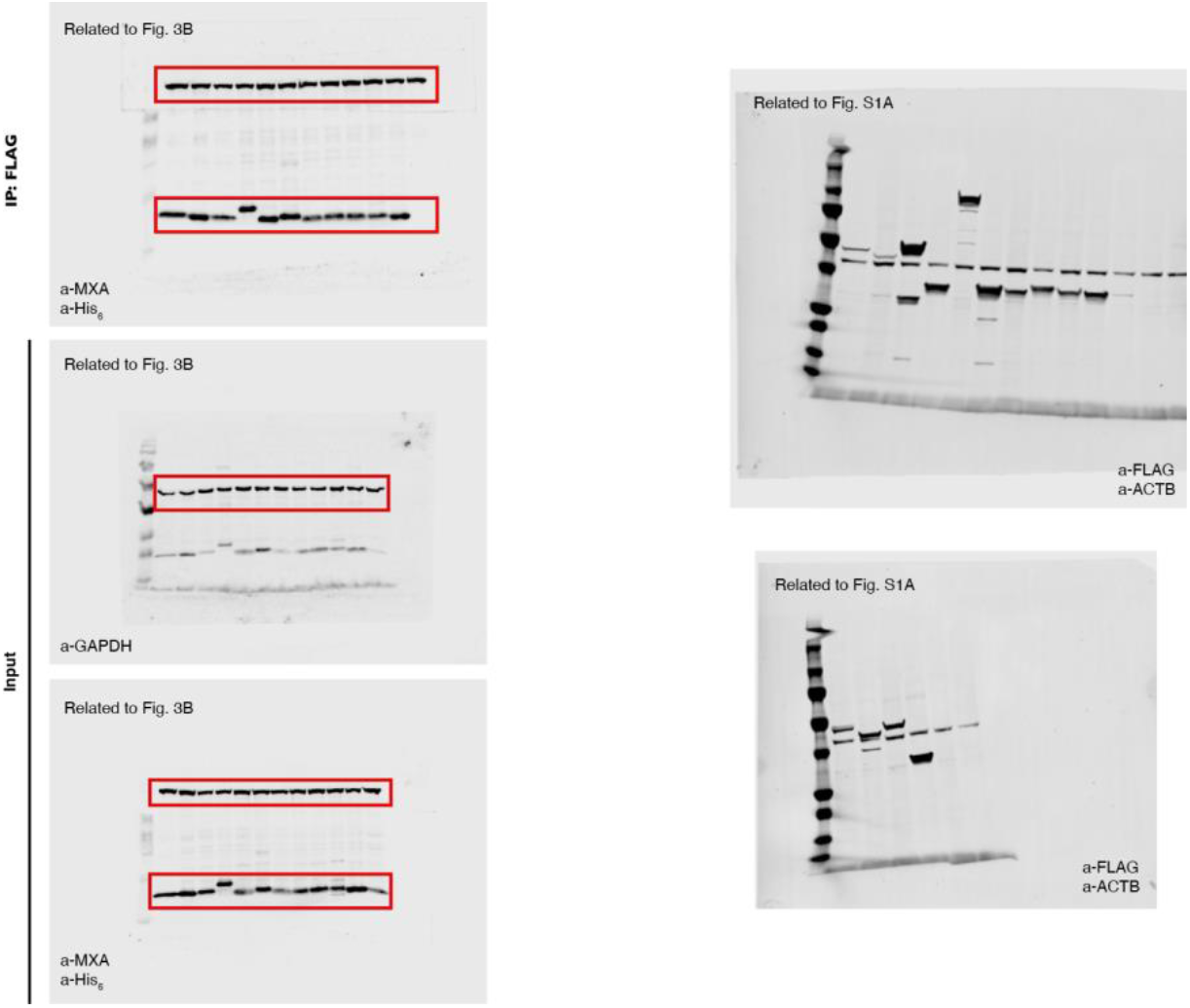
Uncropped immunoblots related to Fig. 3B and Fig. S1A. Antibodies are shown in the blot. For Fig. 3B, the cropped areas in the Figure are highlighted by a red box.

